# Clustered cytoneme interactions contribute to peripheral nervous system patterning in Drosophila

**DOI:** 10.64898/2026.09.11.750707

**Authors:** Sushmita Kundu, Christoph Cunningham, Ginger L. Hunter

## Abstract

The formation of patterned adult tissues requires precise spatiotemporal communication between cells that comprise the tissue during development. Organization of the dorsal thoracic sensory bristles of the fruit fly Drosophila melanogaster relies in part on the activity of cell protrusions called cytonemes. Experimental studies have identified several of the mechanisms that govern individual cytoneme dynamics in the patterning PNS, and theoretical studies have considered the effects of their collective activity. Here, using in vivo and ex vivo imaging strategies combined with fly genetics, we show that cytonemes form clusters along the basal surface and that association with clusters promotes the dynamics of individual cytonemes. Clusters are dynamic and transient organizations of cytonemes, and we show that when their movement is perturbed ex vivo through locally aligned nanofibers, PNS patterning is disrupted. We show evidence that cytoneme interactions via clusters is supported by the cell adhesion molecule E-cadherin, as reducing E-cadherin expression leads to changes in cluster dynamics, ability to promote cytoneme length, and overall patterning of the PNS. Taken together, our results support a model in which complex cytoneme interactions promote the dynamics of individual cytonemes. We propose that the enhanced dynamics and stability provided by increased cell-cell interactions may support both the range and strength of local signaling events.

## Introduction

The spatiotemporal coordination of local cell behaviors and fates is essential for the development and homeostasis of larger scale tissues. A classic example of this is the organization of the peripheral nervous system (PNS) in animals: in order to form a functional system whose components are distributed across the animal body, cells of the nervous system must coordinate the positioning and connectivity of multiple cell types. Cell-cell communication by the secretion of morphogens or through contact during juxtacrine signaling supports local control of patterning, but how these local mechanisms scale across larger tissue regions is still unclear. One cell-based mechanism for long-range signaling is cytonemes. Cytonemes are long (< 100 microns), thin, actin-based cellular protrusions that carry morphogens along their length Liu *et al*. (2026). Cytonemes have been documented in several vertebrate and invertebrate developing tissues Hall *et al*. (2024); Huang *et al*. (2019); Sanders *et al*. (2013); Zhang *et al*. (2024), and many studies show disruption of cytoneme formation or activity leads to perturbations in patterning and tissue function Bischoff *et al*. (2013); Cohen *et al*. (2010). In the PNS, incorrectly positioning or timing the correct fate of a neuron can lead to delays or failures in connectivity.

Like conventional filopodia, cytonemes initiate from the surface of signal sending or receiving cells, and extend. Cytonemes are primarily actin-based structures, similar in many respects to filopodia. Several of the proteins that help form and regulate filopodia are also found to play a role in cytonemes, including Rho family GTPases Rac and Cdc42 Couto *et al*. (2017); Stanganello *et al*. (2015), Diaphanous Roy *et al*. (2014), SCAR Georgiou and Baum (2010), Enabled Sutcliffe *et al*. (2025), Ezrin-Radixin-Moesin (ERM) and regulators De Joussineau *et al*. (2003); Rambaud *et al*. (2025), Myosins Clements *et al*. (2024); Hall *et al*. (2021), and Cofilin Sanders *et al*. (2013). When the activity of positive regulators is perturbed by knockdown or dominant negatives, the formation of cytonemes is reduced (e.g., De Joussineau *et al*. (2003). In addition to extension and retraction activity, cytonemes engage with their environment both through cell-matrix and cell-cell interactions. Support from cell-matrix interactions can support target searching: during larval development, cytoneme-mediated signaling occurs between the wing disc and the air sac primordium Huang and Kornberg (2015); Roy *et al*. (2014). Cell-matrix proteins including Drosophila ß-integrin subunit Myospheroid and its extracellular matrix ligand Laminin are required both for the formation of cytonemes and the ability of formed cytonemes to reach their target tissue in this system Huang and Kornberg (2016). General adhesion between cells via cellular protrusions can involve cell-cell adhesion proteins like cadherins Biswas and Zaidel-Bar (2017); Schwabe *et al*. (2013); Liu *et al*. (2008). Transmembrane proteins Neuroglian and Capricious mediate cell-cell adhesion, notably in the developing nervous system, but also has been shown to play a role in the targeting of cytonemes during air sac primordium development Roy *et al*. (2014). In addition to surface proteins that mediate cell-cell adhesion, interactions between cytonemes can be more generally supported by surface signaling proteins acting as CAMs (cell adhesion molecules). For example, a recent study shows that ligand/receptor interactions between membrane-anchored Branchless (encoding FGF) and Breathless (encoding FGFR) promotes the formation of cytonemes during wing disc development Du *et al*. (2022). Extending this model, any cell surface ligand-receptor pairs could constitute CAMs, and the adhesion strength that they confer on cells may be dependent on both the surface area of contact as well as the lifetime and number of ligand-receptor interactions present.

The patterning epithelium of the fruit fly dorsal thorax is a model system for the study of how cytonemes signal and interact with each other Cohen *et al*. (2010); Hunter *et al*. (2019); Lacoste *et al*. (2022). In adult flies, this region is covered by a repeating spot array of sensory bristles. The wild type spacing of each bristle precursor cell within the epithelium requires long-range Notch signaling supported by cytonemes. Over the course of several hours in pupal stages, long actin-rich cell projections extend from the basal surface of each cell in the bipotential epithelium Clements *et al*. (2024); Cohen *et al*. (2010); Georgiou and Baum (2010); De Joussineau *et al*. (2003); Hunter *et al*. (2019); Renaud and Simpson (2001). Each neuroepithelial cell undergoes a binary fate decision, to remain epithelial or adopt a pro-neural cell fate. This binary decision is facilitated by Notch signaling such that proneural cells repress Notch receptor expression and increase expression of proneural genes, while epithelial cells become Notch activated and repress expression of proneural genes Furman and Bukharina (2008); Lewis (1996); Troost *et al*. (2015). Proneural cells will further divide to generate the cells that produce a functional sensory bristle of the PNS. Both Notch and Delta proteins are observed along the lengths of the cytonemes emanating from the basal surface of the notum epithelium during patterning stages Clements *et al*. (2024); Hunter *et al*. (2019). Mathematical and experimental data indicate that these cytonemes are an important source of signaling for the proper positioning of pro-neural cells in the epithelium Cohen *et al*. (2010); Hadjivasiliou *et al*. (2016). Behavioral experiments show that cytoneme-mediated signaling between notum bristle precursor cells is required for wildtype axonogenesis and subsequently, in adults, functional PNS responses Lacoste *et al*. (2022). Since bristles and the mechanosensory neurons that innervate them are required for several adult behaviors, such as grooming, missing bristles or improperly placed bristles would be expected to lead to failure or delays in responding to environmental stimuli Vandervorst and Ghysen (1980).

Many of the mechanisms discussed above that regulate cytonemes also are observed during long-distance Notch signaling in the patterning notum. Notum cytonemes have been shown to require regulators of the actin cytoskeleton for their activity. Loss of SCAR or Rac activity leads to defects in spacing of proneural cells, consistent with defects in long-range Notch signaling Cohen *et al*. (2010); Georgiou and Baum (2010). However, since the Notch signaling mechanism itself requires wild type actin dynamics in the context of ligand/receptor endocytosis Le Borgne and Schweisguth (2003); Langridge and Struhl (2017), it remains unclear whether long-range Notch phenotypes in actin regulator knockdown tissues are the result of cytoneme defects or endocytosis defects (or some combination of both). The notum epithelium is a single layer of cells, with a thin laminin-based basement membrane directly underneath Mehaffey *et al*. (2024). It is not currently known whether cell-cell or cell-matrix adhesions support notum cytoneme signaling during patterning stages. Finally, there is some evidence that Drosophila Notch receptor and Delta ligand act as CAMs Fehon *et al*. (1990). When Drosophila S2 cells expressing either Notch or Delta are co-cultured, cells aggregate. Canonically, Notch receptor is cleaved and endocytosed upon interaction with Delta in trans, so the overall adhesive stability of Notch-Delta as CAMs may rely on the formation of numerous ligand-receptor interactions across the cell-cell interface.

Despite evidence that contact between cytonemes and their targets is important for signaling and therefore overall patterning, there is a gap in our knowledge about the mechanisms and dynamics that characterize cytoneme-cytoneme interactions. The extent of cytoneme contact with their targets may vary from localized contacts near the tips to extensive contacts along the lengths González-Méndez *et al*. (2017); Huang and Kornberg (2015); Hunter *et al*. (2019); Patel *et al*. (2022); Roy *et al*. (2014), as determined by GRASP-based cell-cell contact reporters Feinberg *et al*. (2008). For signaling paradigms that require contact, like Notch and other juxtacrine signaling systems, signaling strength varies with surface area of contact Shaya *et al*. (2017). This is important in patterning tissues because variability in signal strength can generate different response dynamics and therefore different patterns. For example, epithelial cells that contact a Delta expressing cell by cytonemes alone in the patterning notum experience a slower Notch activation than cells that are adjacent to the Delta source, contacting by both cytonemes and lateral cell-cell surfaceHadjivasiliou *et al*. (2016); Hunter *et al*. (2016). Slower Notch activation has consequences for patterning including extended time in lateral inhibition stages. Understanding how cytonemes dynamically interact with each other during patterning stages may provide some mechanistic insight into how they effectively propagate signals.

In this study, we examine how cytonemes interact with each other in vivo, during a Notch-dependent tissue patterning event, which is the organization of the peripheral nervous system on the dorsal thorax of the developing fruit fly. We show that cytonemes interact with each other in complex ways, like dynamic clusters that move across the basal surface of the tissue. These clusters promote the length and lifetime of individual cytonemes, which suggests positive reinforcement of cytoneme signaling capacity. When blocked ex vivo through locally aligned nanofibers, cytonemes are inhibited in length and dynamics, which leads to patterning defects in the notum. Finally, we show that the cell-cell adhesion protein E-cadherin plays a role in the stability of cytoneme clusters. Together, this work contributes to our understanding of how local signaling events are propagated across larger regions of patterning tissues, as well as our understanding of the proteins that support cytoneme mediated signaling.

## Materials and methods

### Fly husbandry

Flies were maintained on standard fly food in vials or bottles. Fly food was made to order (JazzMix, Fisher Scientific) or purchase pre-made (Lab Express). Fly stocks were kept at 18 °C-25 °C as needed. White prepupae (0 hAP) were collected and aged at 18°C for 24 hours (to 12 hAP). Stocks used are listed in Table 1.

**Table 1.** Fly stocks used in this study.

| Stock | Source<sup>a</sup> |
| --- | --- |
| sGMCA | D.Kiehart Lab |
| senseless-GFP | BDSC 38666 |
| UAS-LifeAct-Ruby/CyO; Pannier-GAL4/TM6B | G.Hunter Lab |
| UAS-Ecadherin-RNAi; UAS-Ftractin-tdTomato/TM6B | G.Hunter Lab |
| Sco/CyO; UAS-Ftractin-tdTomato/TM2 | BDSC 58988 |
| Neur-GMCA/CyO; pannier-GAL4/TM6B | G. Hunter Lab |
| UAS-LifeAct-Ruby | BDSC 35545 |
| UAS-LifeAct-GFP | BDSC 35544 |
| UAS-zyxin-mCherry | BDSC 28875 |
| zyxin-GFP | BDSC 94780 |
| shotgun-GFP | BDSC 60584 |
| UAS-Ecadherin RNAi | BDSC 38207 |
| UAS-zyxin RNAi | BDSC 36716 |
| UAS-LexA RNAi | BDSC 67947 |
| IF/CyO; pannier-GAL4/TM6B | G. Hunter Lab |

<sup>a</sup> BDSC = Bloomington Drosophila Stock Center.**Table 2** Other reagents used in this study

### In vivo microscopy

12 hAP pupae were affixed to microscopy slides using double sided tape and pupal cases were removed using forceps. A #1.5 glass coverslip coated in a thin film of halocarbon oil (1:1 HC27:700)(Sigma H8898 and H8773) was placed on top of spacers to prevent damage to the pupae. Pupae were imaged on either an inverted Leica SP8 line scanning confocal microscope or an inverted Nikon Ti-2 spinning disc confocal microscope at room temperature. LASX software was used to capture timelapse data sets on the Leica confocal. Nikon Elements software was used to capture timelapse data sets on the Nikon confocal.

### Image analysis and statistics

All images were processed and analyzed using FIJI/ImageJ Schindelin *et al*. (2012). For PIV analysis, a FIJI plug in was used Tseng *et al*. (2012). We adjusted window size based on the measured size of cytoneme structures observed in vivo. Python script was implemented to compile the PIV output and determine the mean magnitude of movement for each time point pair. Bristle pattern density in vivo was determined by counting the number of bristles in a 100 × 100 micron ROI in the pannier domain, determined by the expression of UAS-LifeActRuby. Bristle spacing ex vivo was determined by measuring the length between pairs of bristles within (but not across) rows. Data sets were uploaded into GraphPad Prism to generate graphs and to run statistical tests. Sample numbers (n) and details of statistical tests used are listed in each figure legend. Unless otherwise stated, mean ± standard deviation is reported.

### Preparation of nanofiber gel solution for ex vivo experiments

Nanofiber monomers (C16-VVAAEE) were synthesized at the Center for Regenerative Medicine Peptide Core Facility, Northwestern University. 1mg of monomer powder was reconstituted in 150mM NaCl, pH brought to 7.2 with 200mM NaOH, to a final concentration of 1mg/mL. The solution was heated at 80 °C for 30 minutes, then cooled for 2 hours at room temperature. Batches were used within 48 hours and maintained at 4 °C. Solutions were brought to room temperature before use on pupae.

### Ex vivo protocol

12 hAP pupae were pinned to PDMS dissection plates and were dissected from the ventral side to expose the notum in 1X PBS (diluted from 10x; Fisher, BP399-500). For live imaging, nota were cut free at the boundary between the thorax and the abdomen. Nota were transferred to a glass-bottom dish (Mattek, P35G-1.5-14-C). Medium was removed and replaced with 2 µL of thrombin (Roche, 10602400001) solution. 2 µL of fibrinogen (EMD Millipore Corp, 341573) was added to induce the formation of a clot to hold the notum close to the coverslip Loubéry and González-Gaitán (2014). After clotting, the dish was gently filled with clone 8 medium (1x Schneider’s medium (Sigma S0146) supplemented with Insect Medium supplement (Sigma I7267) and FBS (Heat inactivated, Genesee Scientific 25-550H).

For live imaging nota embedded in nanofibers, after nota were transferred to a Mattek dish and excess medium was removed, room temperature nanofiber gel solution was added. Nanofiber polymerization is calcium dependent, and 1X Schneider’s medium contains 5.4 mM CaCl_2_ which is sufficient to induce gelation. Excess fluid was removed, then clot formation was performed and the dish filled with clone 8 medium. Dishes with dissected nota were then gently mounted onto the confocal microscope for imaging.

For short- and long-term patterning experiments with nanofibers, nota were dissected from 12 h AP pupae in 1X PBS, then transferred to room temperature nanofiber gel solution. Each notum was then pipetted gently into a dish containing 1 mL clone 8 medium. We used visual confirmation to ensure nota were embedded in gel (i.e., nota that were successfully embedded could be moved in the dish by gently pushing the gel around it, without touching the tissue itself). Dishes were kept in a humidified chamber for 10-20 minutes (short-term) to 6 hours (to 18 hAP; long-term). At this point, clone 8 medium was removed and replaced with 4% paraformaldehyde solution to fix, followed by immunofluorescence as outlined below.

### Immunofluorescence and stainings

12 hAP pupae were pinned to PDMS (Dow Sylgard 184, Ellsworth) dissection plates and were dissected from the ventral side to expose the notum in 1x PBS. PBS was replaced by 4% paraformaldehyde solution (EMS #15710) in 1X PBS for 20 minutes at room temperature. Tissues were washed in 1x PBST (PBS with 0.1% Triton x-100, Thermoscientific A16046.AP) four times, then blocked in 50% blocking buffer (3% FBS, 5% BSA (Genesee Scientific 25-529) in 1X PBS) for 1 hour. Primary antibody solutions were prepared in 1x PBST with 5% block as listed in table 1 and incubated with the tissue for 1hr (room temperature) to overnight (4°C). Tissues were washed 3 times for 10 minutes each with 1X PBST. Secondary antibody solutions were prepared in 1X PBST, with any fluorescence stains needed, as listed in table Tissues were incubated with secondary solution for 1hr (room temperature) to overnight (4°C). Tissues were washed 3 times for 10 minutes each with 1X PBST. Tissues were equilibrated in 50% glycerol (Thermoscientific J61059.K2) /1x PBS overnight at 4 °C, then mounted on standard glass microscopy slides with #1.5 glass coverslips sealed with nail varnish. Slides were kept in the dark at 4 °C until imaging. For tissues where staining only was performed (no antibodies), dissected nota were directly placed into stain solution prepared in 1X PBST without blocking.

## Results and discussion

### Collective interactions of cytonemes during patterning stages

In order to observe large scale cytoneme interactions during patterning stages (12-18 hours after pupariation, hAP), we use live confocal microscopy to visualize all cytonemes in the central notum epithelium using UAS-LifeAct-Ruby under control of the pannier-GAL4 driver Brand and Perrimon (1993) (Figure 1A). Fluorescently tagged LifeAct binds to and allows visualization of the filamentous actin cytoskeleton Melak *et al*. (2017). Since cytonemes are primarily actin based structures, this allows us to observe them in bulk, however it does not allow us to have single cell resolution since all central notum cells express pannier-GAL4 (pnr-GAL4). Importantly, in this background we cannot distinguish between bipotential neuroepithelial cells that remain epithelial and those that acquire a pro-neural cell fate until 18 hAP. We observed junctional LifeAct-Ruby localization in the apical planes, consistent with epithelial organization (Figure 1A, apical). We also observed cytonemes extending from the basal surface of all epithelial cells (Figure 1A, basal). We were surprised to find that the distribution of cytonemes across the basal surface is not uniform. We observed that there are areas of increased LifeAct fluorescence intensity (Figure 1A, arrows), which we interpret to mean there are more cytonemes per unit area at these locations. These areas of increased LifeAct fluorescence intensity colocalize with increased membrane reporter fluorescence (mCD8-GFP, Figure 1B), which suggests that they are cytonemes, and not intracellular actin structures. To ensure this was not an artifact of expressing LifeAct throughout the tissue, we also separately expressed pnr-GAL4 > UAS-Ftractin-tdTomato and observed the same clustering (Figure S1A). We also tested a GAL4-independent filamentous actin reporter sGMCA (GFP-tagged constitutively active moesin ubiquitously expressed under control of the myosin regulatory light chain promoter)(Edwards et al., 1997). Ubiquitous expression of a GFP-tagged actin reporter increases the background noise from other tissues underneath the notum, but we observe cytoneme clusters in these pupae as well (Figure S1B).

**Figure 1.**
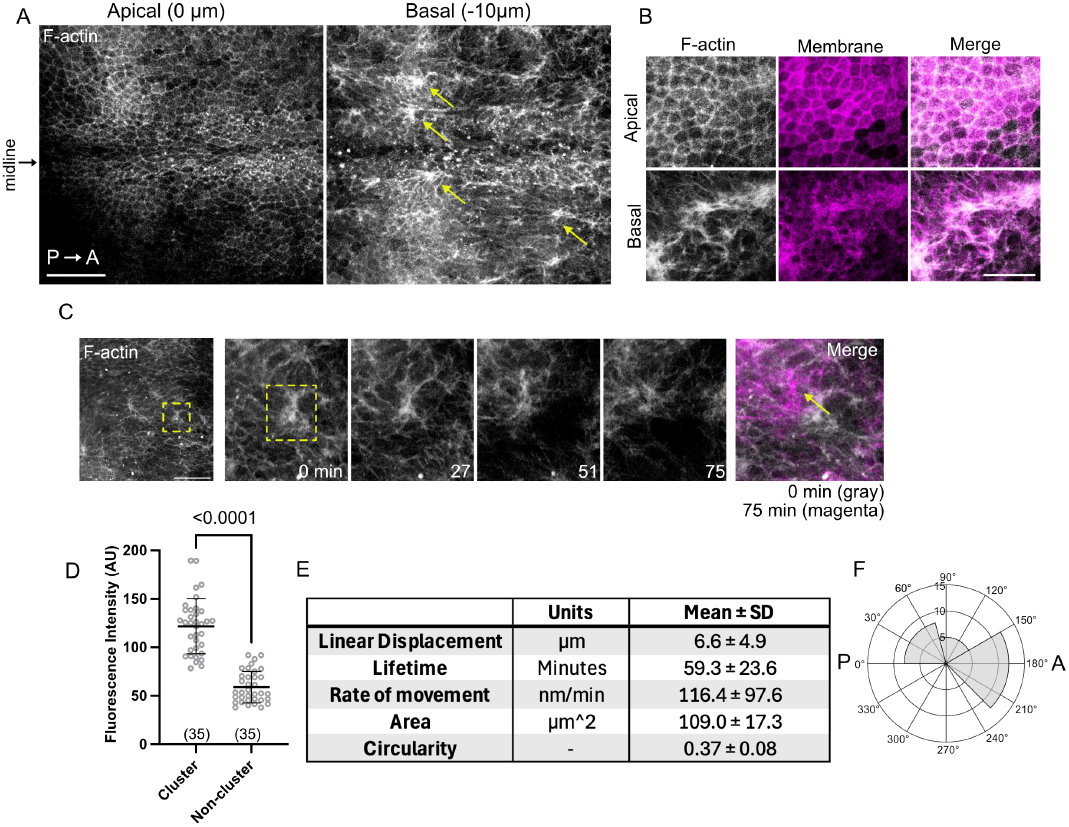
Cytonemes are not uniformly distributed on the basal surface of the patterning. (A) Apical and basal view of a 12 hAP patterning central notum expressing the filamentous actin reporter LifeAct-Ruby under the pannier-GAL4 driver. Animal midine is indicated, and anterior is to the right. Yellow arrows point to clusters of cytonemes on the basal surface. Scale bar, 50 µm. (B) Apical and basal view of a 12 hAP notum expressing LifeAct-Ruby (gray) and the membrane marker mCD8-GFP (magenta) under pannier-GAL4. (C)Timelapse imaging of a cytoneme cluster moving in vivo. Yellow box highlights a cluster. Final merged image overlays the first (gray) and last (magenta) time points to demonstrate the extent of movement. (D) Mean fluorescence intensity on the basal surface for ROIs within and outside of (non) clusters. n = 35 clusters. (E) Summary data for manual analysis of cytoneme clusters in vivo, for n = 21 clusters across 6 pupae. (F) Roseplot indicating the angles of movement for clusters analyzed in (E) aligned to the animal midline (0°). A, anterior. P, posterior.

We also observed that these areas of increased fluorescence intensity are dynamic: they assemble, move across areas of tissue, and disassemble (Figure 1C). We made several manual measurements to describe these dynamics (Figure 1D-F). Our manual measurements use LifeAct fluorescence intensity to distinguish between areas with increased cytoneme densities compared to those with less (Figure 1D). Areas of decreased LifeAct fluorescence intensities still exhibit cytonemes. We find that cytoneme clusters cover large areas relative to both the apical area of an average notum epithelial cell at this stage (diameter 9.4 ± 1.7 µm, n = 50 cells in 5 pupae; 69.4 µm^2^) and the maximum length of individual cytonemes (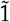0 µm, Cohen *et al*. (2010); Hunter *et al*. (2019). Clusters have relatively long lifetimes relative to the average lifetime of an individual cytoneme (10 minutes, Hunter *et al*. (2019). The rate of movement of the cytoneme cluster is also slower than extension and retraction rates of individual cytonemes (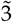0 nm/s, Hunter *et al*. (2019). Finally, we followed the trajectories of movement of the cytoneme structures. We find that they do have persistent movement and that much, but not all, of this movement tends to happen along the anteroposterior axis (Figure 1F). These observations indicate that cytonemes create complex interactions with each other during notum PNS patterning stages. The interaction clusters are limited in space, such that not all cytonemes at a given time are participating in a cluster. From our observations, movement of a cluster comes from both movement as a unit after formation and apparent movement due to individual cytonemes extending into and retracting from the main cluster. Given previous observations that cytoneme interactions across the basal surface are under tension Hunter *et al*. (2019), some of the unit movement may be due to the balance of active contractive forces between interacting cytonemes, and some of the movement may be apparent due to the incorporation or retraction of individual cytonemes, especially at the edges of the cluster.

### Individual cytonemes interact with collective structures

To begin to understand the relationship between the activity of individual cytonemes and the behavior of clusters, we next used a dual-color approach to label individual cells and in a background where cytoneme clusters are also visible. We expressed the filamentous actin reporter GFP-moe^CA^ Edwards *et al*. (1997) under the neuralized promoter (neur-GMCA) in the same background as pnr-GAL4 > UAS-LifeAct-Ruby or UAS-Ftractin-tdTomato. Together this background expresses a red fluorescent actin reporter in all cells in the central notum while also expressing a green fluorescent actin reporter in the pro-neural, bristle precursor cells (Figure 2A). We were surprised to find that the cytoneme clusters are frequently formed along and in-between existing rows at 12-13 hAP. The inbetween space corresponds to the actively patterning rows 2 and 4 Corson *et al*. (2017). Together with the data in Figure 1A and 1F, it appears that clusters form and move parallel to bristle precursor rows during patterning stages. It is important to note that while there are some incidences of apoptosis in the notum during these stages, which lead to extruding cells that can appear to have increased fluorescence intensity via F-actin reporters, we do not observe blebbing or cell delamination in the cytonemes clusters to suggest that these clusters occur concurrently with apoptotic cells.

**Figure 2.**
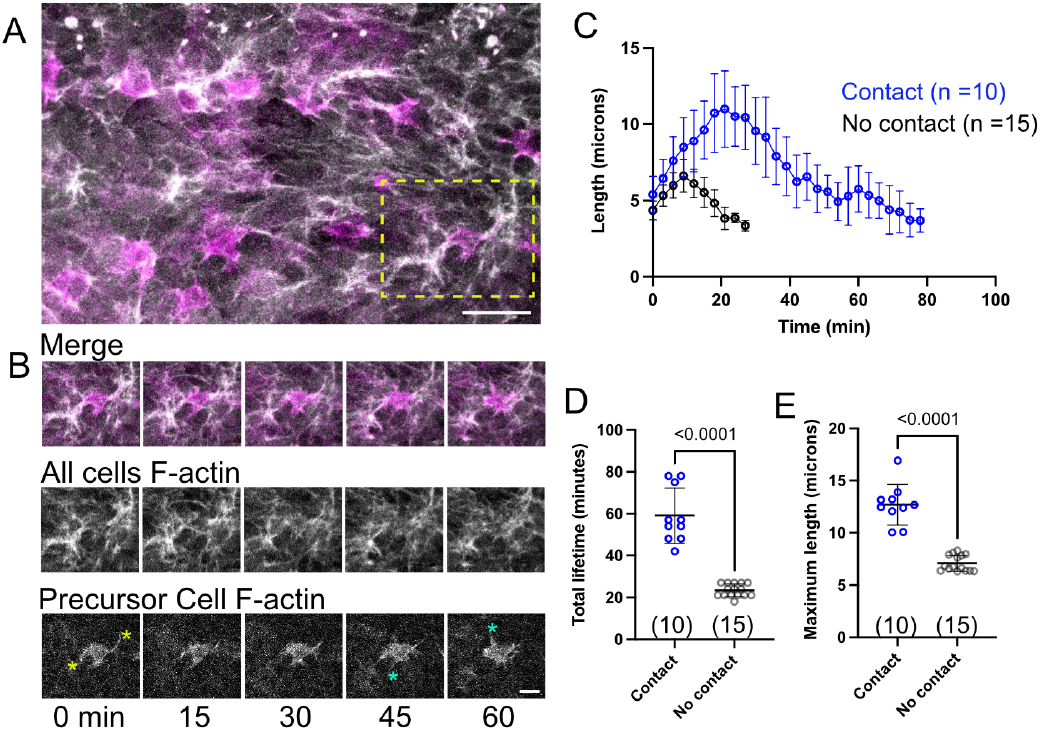
Clustering promotes cytoneme dynamics. (A) Large scale cytoneme organization in tissues expressing LifeAct-Ruby to label filamentous actin in all cells (gray) and GFP-moe^CA^ to label filamentous actin in bristle cell precursors only (magenta). Scale bar 20 µm. (B) Timelapse images of the inset in (A, yellow box). Yellow asterisks label cytonemes that contact clusters, and cyan asterisks label cytonemes that do not contact clusters. Scale bar, 10 µm. (C) Manual tracking of individual bristle cell precursor cytonemes that contact (blue) or do not contact (black) clusters. Mean ± SD shown. n = number of individual filopodia measured, across 4 nota. (D) Total lifetime of cytoneme from initial extension to complete retraction for cells that contact (blue) or do not contact (black) clusters. Mean ± SD shown. Data set is same as (C). (E) Maximum length of cytoneme during its total lifetime for cells that contact (blue) or do not contact (black) clusters. Mean ± SD shown. Data set is same as (C). p-values are determined by student’s t-test.

We observe that bristle precursor cells extend some individual cytonemes towards and into the clusters (Figure 2B). We do not observe that cytonemes from bristle precursor cells preferentially extend towards the clusters. This indicates that the predominant cell type participating in collective cytoneme interactions are those epithelial cells which are actively engaged in lateral inhibition. Although bristle precursor cytonemes do not preferentially extend towards cytoneme clusters, we do observe that the ones that do have an extended lifetime compared to those that extend in other directions (Figure 2C and D). We also observe that cytonemes that extend towards and into clusters have an increased maximum length relative to those that extend in other directions (Figure 2C and E). Therefore interactions with clusters are positively enforcing the dynamics of existing cytonemes, without necessarily promoting their formation. Cytoneme clusters could represent interactions between cell bodies and cellular projections from numerous nearby differentiating cells. Since Notch and Delta are surface signaling proteins found on the cytonemes, we propose that cytoneme clusters represent signaling hotspots under two potential models. First, clusters increase the number of cells that an individual cytoneme can come into contact with relative to cytonemes that do not extend into clusters; second, clusters provide stability, as increased lifetime could increase the probability of successful signaling events between contacting cytonemes. However, from this study it is not clear yet whether clusters of cytonemes are actively formed by cells in the tissues or are an emergent property of cytoneme interactions. Furthermore, due to limitations in our available GAL4 drivers, we do not know if this effect on cytoneme length and lifetime is specific to cytonemes on bristle cell precursors, or for any cytoneme that interacts with a cluster.

### Basal cytoneme dynamics promote wild type bristle patterning

To determine whether cytoneme clusters play any role in supporting cell-cell signaling during pattern formation, their formation or dynamics need to be perturbed and the consequences on patterning quantified. However, there are no known genetic targets in Drosophila whose knockdown can specifically block the formation of cytonemes without also affecting actin polymerization elsewhere inside the cell (e.g., SCAR, Rac, Cohen *et al*. (2010); Georgiou and Baum (2010). To overcome this problem, we developed an ex vivo approach and embedded dissected nota in peptide amphiphile nanofiber gels Berns *et al*. (2014). We hypothesized that these nanofiber gels could be polymerized around the dissected nota to provide physical ‘road blocks’ to cytoneme activity (Figure 3A). Our protocol for polymerizing the nanofibers does not generate global alignment across the tissue (i.e., we do not apply a pulling force on the gels as they form), but we do expect that there will be local regions of nanofiber alignment. Importantly, we do not know if or how cells are interacting with the nanofibers, other than as physical barriers as previously shown for cells in culture Berns *et al*. (2014). When we incubate nota in nanofiber gels for short periods (< 20 minutes), followed by fixation and phalloidin staining, we observe that cytoneme connectivity decreases, in part due to apparent reduction in cytoneme length compared to unembedded control tissues (Figure 3B). Importantly, at these time scales, we do not observe any evidence of increased cell death or mitosis (observed by nuclear morphology and DAPI stain), both of which may be associated with retraction of cytonemes Cohen *et al*. (2010); Rosa *et al*. (2015).

**Figure 3.**
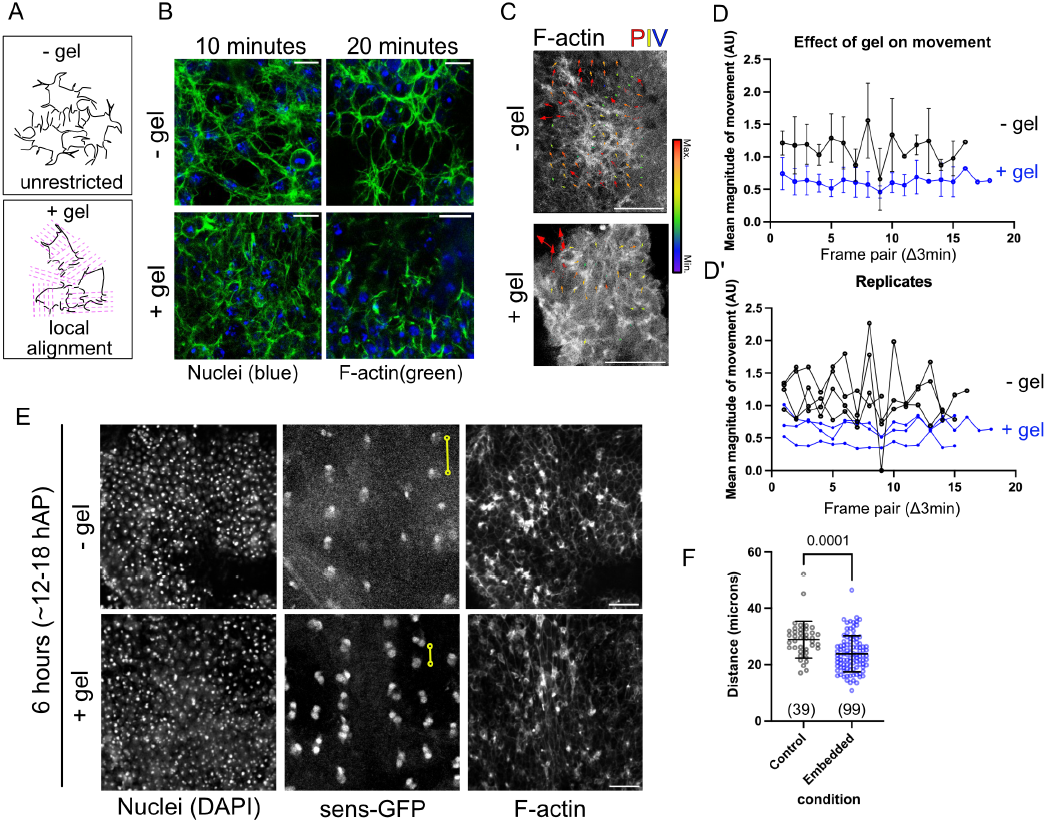
Nanofiber gels disrupt basal cell morphology and dynamics. (A) Schematic of the effects of nanofiber gel (+gel, pink lines) treatment relative to control (−gel). (B) Short term incubation of dissected 13 h AP nota with (+gel) and without (−gel) embedding in nanofiber gels. After the time period indicated, tissues were fixed and stained with phalloidin to visualize filamentous actin (F-actin) and DAPI (nuclei). Scale bars, 10 µm. (C) Live imaging of dissected 12 h AP nota with (+gel) and without (−gel) embedding in nanofiber gels. Nota express a constitutive filamentous actin marker tagged with GFP (sGMCA). PIV overlay display the movement observed between two frames. (D-D’) Mean magnitude of movement per frame pair using PIV analysis. Vector size and color indicate magnitude (red, big = large movement; blue/violet, small = small movement). (D)shows mean ± SD summary plot and (D’) shows all replicates. (E)Long term incubation of dissected 12 h AP nota with (+gel) and without (−gel) embedding in nanofiber gels. Nota express GFP tagged senseless, which is expressed in pro-neural cells. After 6 hours, tissues were fixed and stained with anti-GFP, phalloidin to visualize filamentous actin (F-actin), and DAPI (nuclei). Scale bar 25 µm. (F) Distance between GFP+ nuclei pairs along rows in control (grey) or nanofiber gel embedded (blue) nota. Yellow lines in (E) indicate how the measurements for (F) were made. Control, n = 39 pairs measured across 3 nota. Embedded, n = 99 pairs measured across 6 nota. P = 0.0001 by Student’s t-test, mean ± SD shown.

Next we asked whether embedding dissected nota in nanofiber gels also reduces the dynamic collective movement of cytonemes. We performed live confocal imaging of dissected tissues expressing sGMCA to visualize the filamentous actin cytoskeleton in the presence and absence of nanofiber gels, and analyzed collective cytoneme movement by particle image velocimetry (PIV) Tseng *et al*. (2012). Although this strategy can capture all basal movement, we used our measurement of the average area of a cluster (Figure 1E) to set the analysis window size for analysis. This approach will filter out some of the movements associated with individual cytonemes which occur at a smaller scales. The ex vivo imaging of nota only allows us to visualize local areas of the tissue, rather than the whole tissue due to tissue folding that can occur during preparation. We find that the presence of nanofiber gels reduces overall movement on the scale of cytoneme networks on the basal surface relative to unembedded controls (Fig 3D-D’). Together with the previous result that the nanofiber gel disrupts cytoneme connectivity, this suggests that our embedding protocol is a good tool for the manipulation of cytonemes ex vivo. However, this tool remains non-specific, and we do not know if the decreased movement we observe by PIV is a defect in creating clusters due to the presence of shorter or fewer cytonemes or a direct disruption to the movement of existing clusters.

Finally, we asked whether nanofiber mediated disruption of cytoneme activity affects overall tissue patterning. Adapting previously published methods for short-term incubation of notum tissues Hunter *et al*. (2019); Loubéry and González-Gaitán (2014), we dissected 12h AP notum epithelia endogenously expressing senseless-GFP (sens-GFP) to label the bristle precursor cell lineage. We then allowed the tissues to pattern to completion in the presence or absence of nanofiber gel. The positioning of notum precursor cells within rows typically completes by 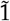8-20 hAP in vivo Cohen *et al*. (2010); Hunter *et al*. (2016). We then fixed the tissues and stained for GFP signal to visualize bristle lineage, filamentous actin to visualize cytonemes, and all nuclei. Nota that are not embedded in nanofiber gels pattern with well-spaced sens-GFP positive bristle cell nuclei in rows (Figure 3E, top), similar to tissues patterning in vivo. Importantly, we observe that both embedded and control bristle precursor cells undergo their characteristic cell divisions, resulting in two or more sens-GFP positive cells clustered together (Figure 3E). The first of these divisions occurs 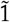6-17 hAP in vivo Roegiers *et al*. (2001); Hunter *et al*. (2016); Lacoste *et al*. (2022). We find that nota embedded in nanofiber gels are able to develop nearly as wild type, with well-spaced sens-GFP positive bristle precursor daughter cells throughout the notum (Figure 3E, bottom). However, when we measure the pairwise distance between GFP positive nuclei in rows, we observe that there is reduced spacing between bristle precursors in embedded nota (Figure 3F). In both control and embedded tissues, we monitored nuclear morphology and do not observe evidence of increased cell death in the central notum. We do observe some necrosis at the cut edges of the dissected tissues. Together with our evidence that short-term nanofiber treatment leads to disrupted cytoneme formation and movement, this result supports a role for cytonemes in the spacing of bristle precursor cells. This finding is consistent with previously performed work that shows when cytoneme formation is reduced by genetic perturbation of the actin regulators SCAR or rac, spacing between bristle precursor cells decreases Cohen *et al*. (2010); Georgiou and Baum (2010).

### Localization of adhesion proteins to cytonemes

Since cytoneme activity and cluster dynamics are required for patterning, we next investigated the molecular mechanisms that allow cytonemes to interact with each other. Based on research showing that filopodia and cytonemes can carry adhesion proteins Jacquemet *et al*. (2019); Junyent *et al*. (2021), we initially hypothesized that either cell-cell adhesion, cell-matrix adhesion, or a combination of both would be essential for the formation of cytoneme clusters. We began by asking which adhesion complexes could be found along the length of notum cytonemes. First, we used the E-cadherin-GFP knock-in line that expresses a GFP-tagged full length E-cadherin under its endogenous promoter to visualize cell-cell adhesion complexes Huang *et al*. (2009). As reported previously Curran *et al*. (2017); Renaud and Simpson (2001), E-cadherin predominantly localizes to the apical adherens junctions of the notum epithelial cells (Figure 4A). We observe that E-cadherin-GFP localizes to puncta along the lengths of cytonemes when co-expressed with LifeAct-Ruby to visualize filamentous actin (Figure 4A). E-cadherin GFP puncta localize to cytoneme clusters as well (Figure 4A). We observe bright GFP puncta localizing to large clusters of cytonemes and smaller clusters as well. E-cadherin is found at low levels across the basal surface, rather than exclusively recruited in cytoneme clusters.

**Figure 4.**
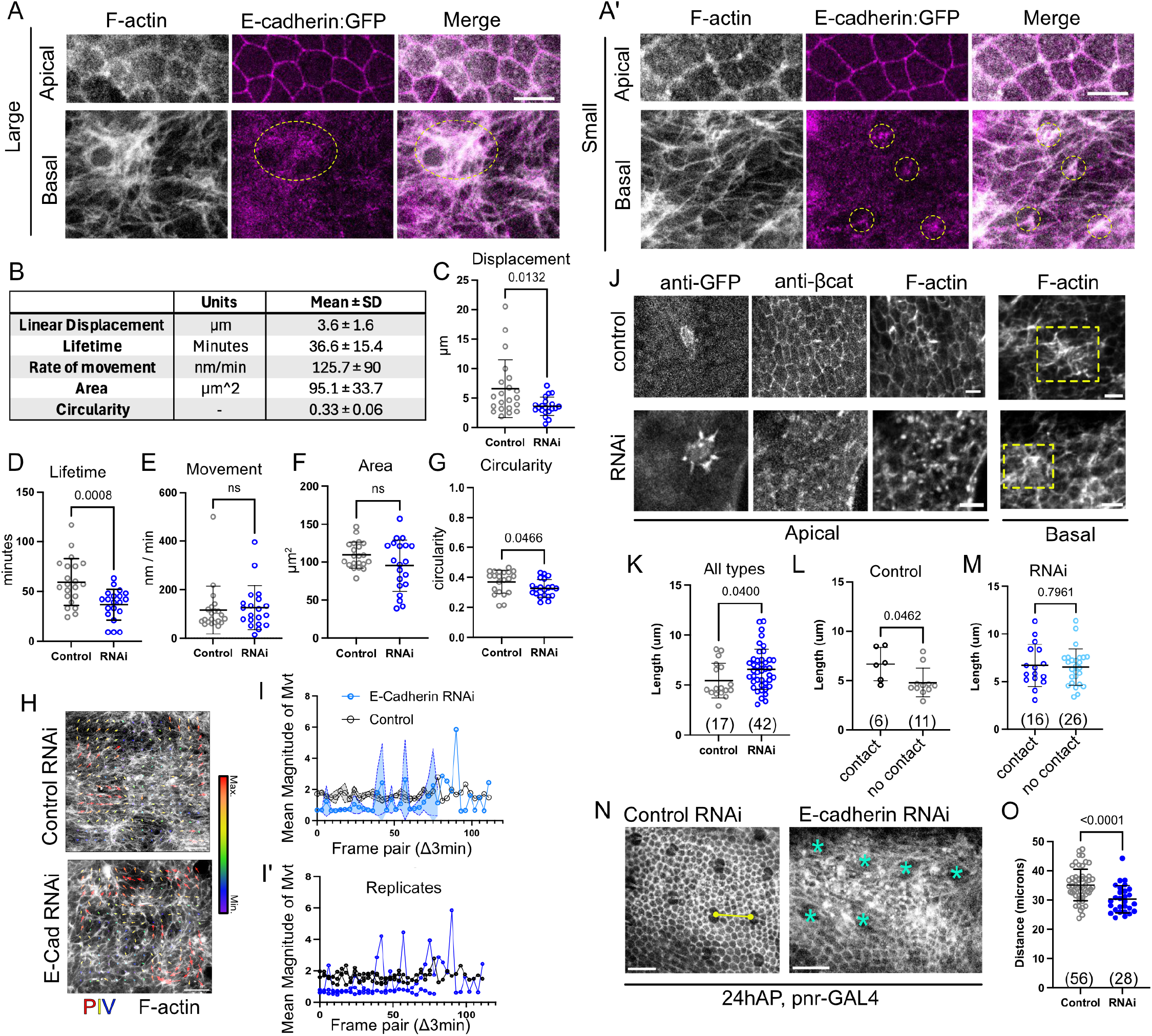
E-cadherin is required for cluster dynamics in vivo. (A-A’) Nota co-expressing the filamentous actin reporter Life-Act Ruby (gray) under the pnr-GAL4 driver with endogenously tagged E-cadherin-GFP (magenta). E-cadherin-GFP localizes apically to cell-cell junctions. E-cadherin-GFP localizes to puncta on the basal surface at both large (A) and small (A’) cytoneme interactions (yellow circles). Scale bars, 10 µm. (B) Summary table of manual tracking of cytoneme clusters in E-cadherin RNAi expressing tissues. N = 20 clusters measured across 6 pupae. (C-G) Graphical representation of data in tables 4B and 1E, using student’s t-test to compare means. Ns, not significant. Mean ± SD shown. (H) PIV overlay displaying the movement observed between two frames of either control RNAi or E-cadherin RNAi expressing nota co-expressing Life-Act Ruby (grey). Vector size and color indicate magnitude (red, big = large movement; blue/violet, small = small movement). (I-I’) PIV analysis of control (black) and E-Cadherin RNAi (blue) cytoneme clusters in vivo. For each frame pair, the mean vector magnitude for the ROI was found and reported. Y-intercept value indicates how large movements are (zero = no cluster movement) and slope of the line indicates whether movement is constant (m = 0) or increasing (positive) or decreasing (negative). Mean ± SEM shown in (I), replicates shown in (I’). (J) Fixed and stained control (top) or E-cadherin RNAi expressing notum (lower) at apical (left) or basal (right) surfaces. Bristle precursor cells express GFP to label their F-actin cytoskeleton. (K) Cytonemes from bristle precursor cells in E-cadherin RNAi expressing tissues exhibit longer cytonemes than control. (L) In control nota, cytonemes that contact clusters are longer than those that do not have contact. (M) In RNAi expressing tissues, cluster contact has no effect on cytoneme length. (N) 24 hAP pupae expressing E-cadherin RNAi showing final bristle cell organization (asterisks). Scale bar, 25 µm. (O) Distance between adjacent bristle cells along rows in 24 hAP nota expressing control RNAi (LexA RNAi, black; n =56) or E-cadherin RNAi (blue; n = 28). P value determined by student’s t-test, mean ± SD shown.

We used a zyxin-GFP endogenous knock-in line to visualized cell-matrix adhesion complexes Singh *et al*. (2026); Stinchfield *et al*. (2024); Yoshigi *et al*. (2005). We did not observe any fluorescence along basal cytonemes (data not shown). We also used a UAS-zyxin-mCherry overexpression line. Overexpression of zyxin-mCherry led to the formation of intracellular aggregates, however a low level of localization was observed to adherens junctions (Figure S4A, arrows). No obvious localization was observed along the lengths of cytonemes, visualized by LifeAct-GFP.

### E-cadherin is required for cytoneme dynamics

We next examined whether E-cadherin is required for cytoneme dynamics and interactions. E-cadherin is an essential protein for maintaining epithelial integrity from early development, and E-cadherin null mutants are embryonic lethal Uemura *et al*. (1996). To study the role of E-cadherin during pupal stages, we expressed UAS-E-cadherin RNAi under the pnr-GAL4 driver at 18 °C, which lowers the efficacy of the GAL4/UAS system Duffy (2002). We considered a pupae to be in a weak E-cadherin knockdown state when we observed filamentous actin disorganization at the apical surface, consistent with published effects of E-cadherin knockdown in the notum Curran *et al*. (2017). In pupae with weak E-cadherin knockdown, we still observe individual cytoneme formation and dynamic behaviors (Figure 4H, J, N). We do not observe any basal actin phenotypes in mild E-cadherin RNAi knockdown nota relative to RNAi controls (UAS-LexA-RNAi; Figure 4H).

We next quantified the dynamics of cytoneme clusters in pupae expressing E-cadherin RNAi. First we analyzed the clusters manually, using fluorescence intensity as a guide, similar to wild type animals (Figure 1). We observe that cytoneme clusters are able to form and are dynamic in pupae expressing E-cadherin RNAi (Figure 4B-G). We find that reduced E-cadherin expression is associated with decreased displacement of clusters in the tissue (Figure 4C), and decreased overall lifetime of the clusters (Figure 4D). We do not see any significant difference in rate of movement (Figure 4E), area (Figure 4F), or circularity (Figure 4G). We do note that although mean area measurements are not different between control RNAi and E-cadherin RNAi clusters, there is increased variability in cluster area when E-cadherin levels are decreased (Figure 4D). Specifically, E-cadherin knockdown tissues exhibit more clusters that are smaller than those found in wildtype, suggesting that the formation of cytoneme clusters is delayed in E-cadherin RNAi tissues.

We also quantified total cytoneme cluster movement in vivo by PIV, as in our ex vivo experiments. We observe that there is reduced basal movement in UAS-E-cadherin RNAi expressing tissues relative to RNAi controls (Figure 4H-I). Compared to our manual analysis (Figure 4B), PIV does not specifically quantify cytoneme clusters, but rather agnostically measures the movement of objects in a window set at the mean size of a cluster. Therefore, we expect that our PIV strategy may pick up additional movements than just that of clusters. We find that decreased E-cadherin expression does not completely stop all basal movement, and that the movement we observe is more noisy than controls (Figure 4I). Finally, we observe no difference from controls in UAS-zyxin RNAi expressing nota (Figure S4B). Together this suggests that robust E-cadherin expression is required for collective cytoneme dynamics.

To understand whether reduced E-cadherin expression leads to changes in how individual cytonemes interact with collective structures, we repeated our dual color experiment (Figure 2) while expressing E-cadherin RNAi throughout the central notum. When we attempted live confocal imaging of the system, we discovered that co-expression of E-cadherin RNAi led to reduced GFP-reporter expression under the neuralized promoter (neur-GMCA) compared to controls. Given previous evidence that robust adherens junctions are required for the apical localization of Notch and Delta proteins Sasaki *et al*. (2007), it is possible that our mild expression of E-cadherin RNAi throughout the notum disrupts lateral inhibition to the extent that additional, activated Notch in bristle precursors leads to the repression of neuralized promoter Miller *et al*. (2014)and decreased GMCA expression. Regardless of the mechanism, we were unable to clearly resolve individual cytonemes extending from the basal surface of bristle precursor cells (as in Figure 2B). As an alternative, we fixed and stained knockdown and control tissues. Under these conditions we can better label and resolve individual cytonemes as well as observe clustering (Figure 4J), even with weakly expressing GMCA cells. Consistent with live imaging of E-cadherin RNAi tissues, we observe disruption of apical F-actin organization in the epithelia. We co-stained for beta-catenin (Figure 4J, middle panel) and also observed failure of beta-catenin to localize strongly to the cell junctions, which is associated with decreased E-cadherin expression (Curran et al., 2017). Since we could not quantify dynamic cytoneme behaviors in live tissues, we investigated the length of cytonemes that were or were not in contact with nearby clusters. Cytoneme clusters were present on the basal surface of both control and E-cadherin RNAi tissues (Figure 4J, basal), visualized by phalloidin staining. When we pool data from all cytonemes measured, we observe that the mean length of observable cytonemes is slightly longer in E-cadherin RNAi expressing nota (6.6 ± 2.0 µm, for 42 cytonemes across 35 cells in 10 pupae; mean ± SD) compared to controls (5.5 ± 1.7 µm, for 17 cytonemes across 14 cells in 4 pupae) (Figure 4K). We next compared individual cytonemes based on whether they were contacting a nearby cytoneme cluster. For controls, we observe that individual cytonemes that contact clusters are longer on average (6.7 ± 1.7 µm, for 6 cytonemes across 5 cells in 2 pupae) compared to individual cytonemes that do not contact clusters (4.8 ± 1.5 µm, for 11 cytonemes across 11 cells in 4 pupae) (Figure 4L). This result is consistent with our in vivo data from Figure 2E, with the caveat that the data in Figure 4 is from fixed tissues and therefore we cannot know what stage of cytoneme life cycle we are measuring (extension, maximum length, or retraction). When E-cadherin expression is decreased, we do not observe a measurable difference in length between cytonemes that contact clusters (6.7 ± 2.2 µm, for 16 cytonemes across 14 cells in 6 pupae) and those that do not (6.5 ± 1.9 µm, for 26 cytonemes across 23 cells in 10 pupae) (Figure 4M).

The results of this experiment demonstrates that, even when not considering the dynamic life cycle of a cytoneme, E-cadherin is required for the ability of cytoneme clusters to promote interactions with individual cytonemes.

Finally, if E-cadherin helps stabilize cytoneme-mediated cell-cell interactions on the basal surface where Notch signaling occurs, we would expect reducing E-cadherin protein levels could reduce the strength of Notch signaling and lead to patterning defects in the notum. Specifically, when long-distance Notch signaling is perturbed, bristle precursor cells are positioned closer together than in wildtype nota Cohen *et al*. (2010); Hadjivasiliou *et al*. (2016). To determine if E-cadherin is supporting long-distance Notch signaling, we quantified the final spacing of sensory bristles at 24 hAP under E-cadherin or control RNAi conditions (Figure 4N-O). We find that reduced E-cadherin expression is associated with reduced spacing between bristles along rows (Figure 4O). This result is consistent with previous observations disrupting long-range Notch signaling via disruption of cytoskeleton components. When we performed a similar experiment to observe the effect of reducing Zyxin expression in the notum, we did not observe any effect on within row bristle spacing by 24 hAP (Figure S2C-D). Since we observe decreased bristle spacing, but not adjacent pairs of bristles, our data suggests that the patterning defect associated with decreased E-cadherin expression is primarily due to inability of cells to propagate long distance Notch signaling. Interestingly, previous experiments in the patterning notum show that over-expression of a dominant negative E-cadherin (pannier-GAL4>UAS-CADH^intra3^) disrupts bristle patterning Renaud and Simpson (2001, 2002): CADH^intra3^ expression in wild type background decreased bristle density - in contrast with our results (Figure 4O). However, in a *scabrous* mutant background, CADH^intra3^ increases bristle density. It is not immediately clear why expression of E-cadherin RNAi or dominant negative E-cadherin should have different bristle density outcomes. It was also observed that CADH^intra3^ expression in the notum disorganized bristle row alignment Renaud and Simpson (2002). Finally, expression of CADH^intra3^ in bristle precursor cells alone was insufficient to disrupt patterning. Importantly, adjacent pairs of bristles were not reported as a phenotype in CADH^intra3^ experiments, indicating successful lateral inhibition between adjacent cells. Our data suggests a model of cooperation between E-cadherin and Notch signaling where cell-cell adhesion mediated by E-cadherin helps to stabilize interactions between cytonemes. Stabilization includes promoting the length and lifetime of individual cytonemes. Since Notch signaling is a contact mediated pathway and Notch receptors and Delta ligands are found along the surface of cytonemes in the notum Hunter *et al*. (2019); Clements *et al*. (2024), stabilized interactions between cytonemes may support effective Notch-Delta interactions. The mechanism by which E-cadherin and Notch cooperate to support signaling in the notum are likely tissue and morphogen specific, as there are other examples of cytoneme signaling in Drosophila that do not exhibit E-cadherin (e.g., Hedgehog signaling during oogenesis Rojas-Rios *et al*. (2012)).

## Conclusions

Here we investigated the how cytonemes interact with each other during the patterning of the fly peripheral nervous system. Our data suggests that cytonemes cluster across the basal surface with the help of E-cadherin mediated cell-cell adhesion, and that interactions within clusters enhances individual cytoneme dynamics, promoting efficient Notch signaling. There is evidence that Drosophila E-cadherin is important for targeting Notch trafficking to apical cell-cell junctions Benhra *et al*. (2011) and evidence for physical interactions between E-cadherin and the Notch intracellular domain at apical cell-cell junctions Sasaki *et al*. (2007). Furthermore, Scabrous, a secreted protein that directly binds to and stabilizes cell surface Notch protein Petruccelli *et al*. (2018); Powell *et al*. (2001)also modifies cell-cell adhesion in the notum Renaud and Simpson (2001). Our results do not rule out a either a model by which E-cadherin could directly support cytoneme-mediated Notch signaling through protein-protein interactions, or an indirect model by which Scabrous participates in modifying both Notch and E-cadherin basally. Since there are many examples of cytonemes interacting with other cytonemes during tissue development, we expect that our observations may be relevant to other settings where complex cell-cell interactions impact the range and strength of local signaling events.

## Acknowledgments

GH conceived of the study. SK, CC, and GH performed the experiments and analysis in this study. SK and GH wrote the manuscript.

We acknowledge our colleague Dr. Shantanu Sur (Clarkson University) for his advice and guidance on the use of PA nanofibers. GH acquired the funding sources for this project. We thank members of the Hunter Lab for their critical reading of this manuscript. We acknowledge use of FlyBase (release FB2026-02 and prior years) to find information on phenotypes and stocks. Stocks obtained from the Bloomington Drosophila Stock Center (NIH P40OD018537) were used in this study (Table 1). Antibodies obtained from the Developmental Studies Hybridoma Bank (DSHB, University of Iowa) were used in this study (Table 2).

**Table 2.**
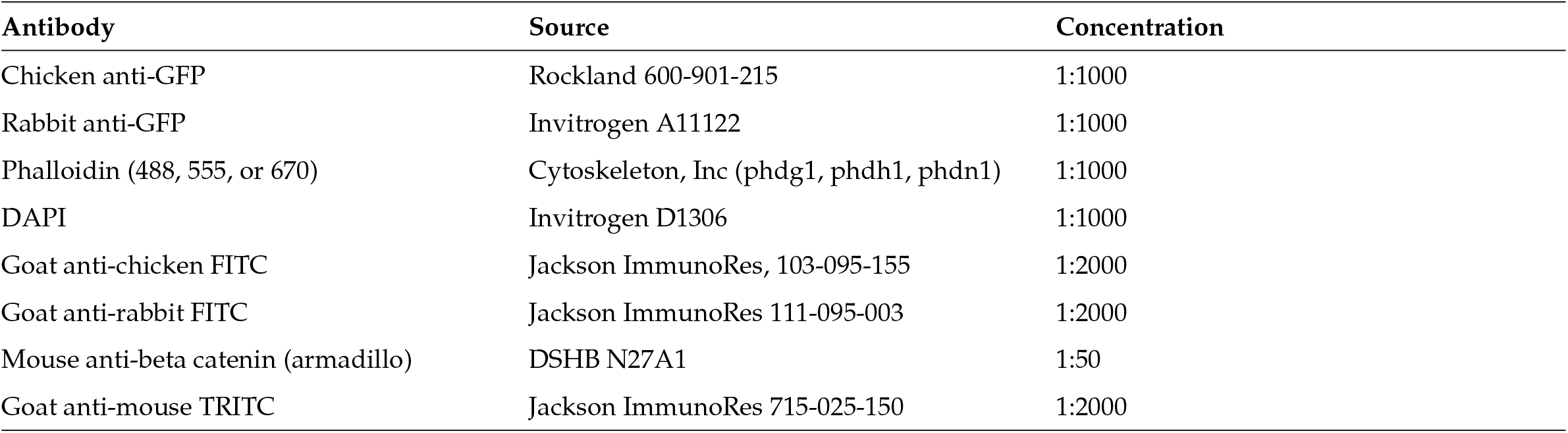
Other reagents used in this study.

| Antibody | Source | Concentration |
| --- | --- | --- |
| Chicken anti-GFP | Rockland 600-901-215 | 1:1000 |
| Rabbit anti-GFP | Invitrogen A11122 | 1:1000 |
| Phalloidin (488, 555, or 670) | Cytoskeleton, Inc (phdg1, phdh1, phdn1) | 1:1000 |
| DAPI | Invitrogen D1306 | 1:1000 |
| Goat anti-chicken FITC | Jackson ImmunoRes, 103-095-155 | 1:2000 |
| Goat anti-rabbit FITC | Jackson ImmunoRes 111-095-003 | 1:2000 |
| Mouse anti-beta catenin (armadillo) | DSHB N27A1 | 1:50 |
| Goat anti-mouse TRITC | Jackson ImmunoRes 715-025-150 | 1:2000 |

---

**Figure 5 Figure S1, related to Fig.1.**
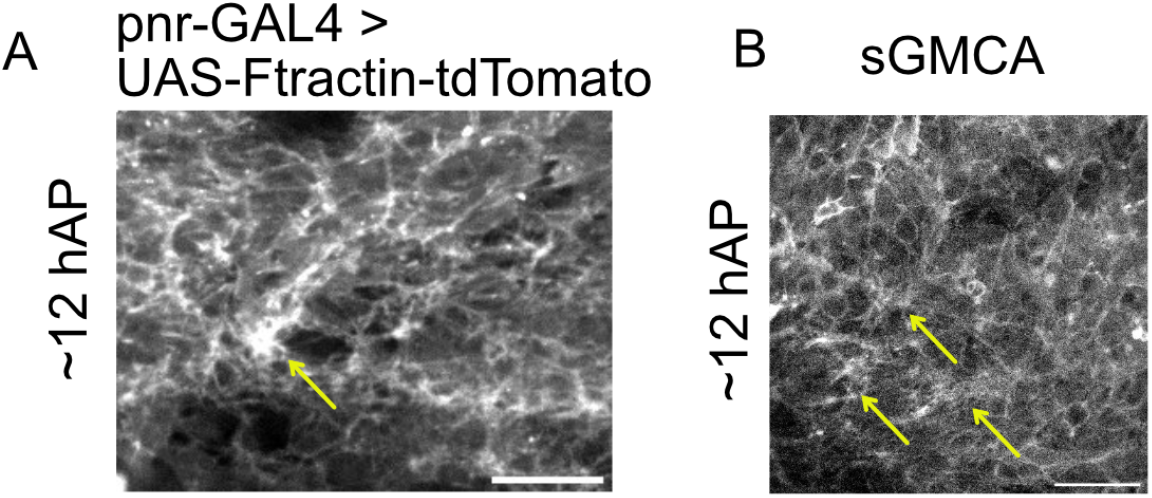
(A) 12 hAP pupae expressing UAS-Ftractin-tdTomato under the pnr-GAL4 driver. (B) 12 hAP pupae expressing sGMCA. Scale bars, 25 µm. Arrows point to cytoneme clusters.

**Figure 6 Figure S2, related to Fig.4.**
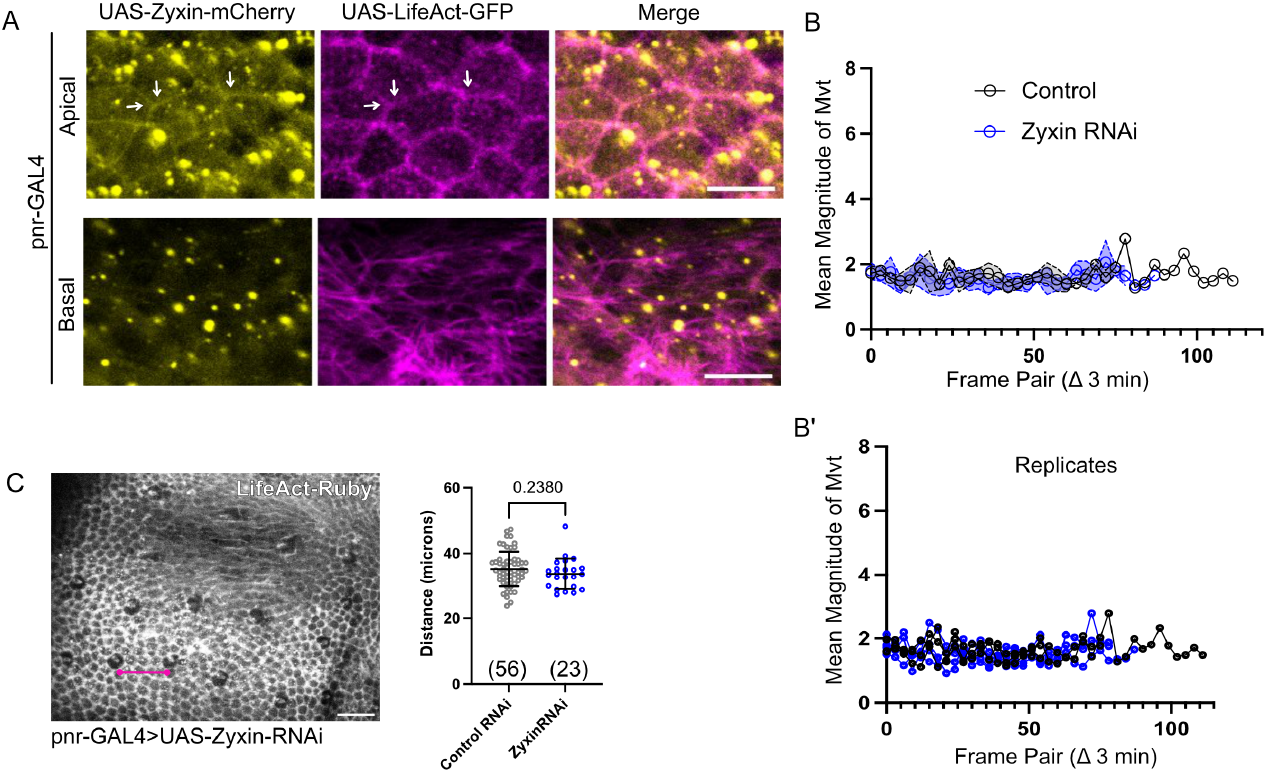
(A) Overexpression of zyxin-mCherry in the pannier domain by pnr-GAL4 at 13 hAP. Apical and basal z-sections shown. Arrows point to apical cell-cell junctions containing both Life-Act and zyxin markers. Scale bar, 10 µm. (B-B’) PIV analysis of basal cytoneme movement in the presence (control, black n = 3 nota) or absence (RNAi, blue n = 4 nota) of zyxin expression. Mean ± SEM shown in (B), replicates shown in (B’). (C) Loss of zyxin expression (blue; n = 23) in the central notum by pnr-GAL4 does not affect the spacing of bristles within rows compared to controls (black; n = 56). Mean ± SD shown. P-value determined by student’s t-test.

## Literature cited

Benhra N, Lallet S, Cotton M, Le Bras S, Dussert A et al. 2011. Ap-1 controls the trafficking of notch and sanpodo toward e-cadherin junctions in sensory organ precursors. Current Biology. 21:87–95.

Berns EJ, Sur S, Pan L, Goldberger JE, Suresh S et al. 2014. Aligned neurite outgrowth and directed cell migration in self-assembled monodomain gels. Biomaterials. 35:185–195.

Bischoff M, Gradilla AC, Seijo I, Andrés G, Rodríguez-Navas C et al. 2013. Cytonemes are required for the establishment of a normal hedgehog morphogen gradient in drosophila epithelia. Nature cell biology. 15:1269–1281.

Biswas KH, Zaidel-Bar R. 2017. Early events in the assembly of e-cadherin adhesions. Experimental cell research. 358:14–19.

Brand AH, Perrimon N. 1993. Targeted gene expression as a means of altering cell fates and generating dominant phenotypes. development. 118:401–415.

Clements R, Smith T, Cowart L, Zhumi J, Sherrod A et al. 2024. Myosin xv is a negative regulator of signaling filopodia during long-range lateral inhibition. Developmental biology. 505:110–121.

Cohen M, Georgiou M, Stevenson NL, Miodownik M, Baum B. 2010. Dynamic filopodia transmit intermittent delta-notch signaling to drive pattern refinement during lateral inhibition. Developmental cell. 19:78–89.

Corson F, Couturier L, Rouault H, Mazouni K, Schweisguth F. 2017. Self-organized notch dynamics generate stereotyped sensory organ patterns in drosophila. Science. 356:eaai7407.

Couto A, Mack NA, Favia L, Georgiou M. 2017. An apicobasal gradient of rac activity determines protrusion form and position. Nature communications. 8:15385.

Curran S, Strandkvist C, Bathmann J, De Gennes M, Kabla A et al. 2017. Myosin ii controls junction fluctuations to guide epithelial tissue ordering. Developmental cell. 43:480–492.

De Joussineau C, Soulé J, Martin M, Anguille C, Montcourrier P et al. 2003. Delta-promoted filopodia mediate long-range lateral inhibition in drosophila. Nature. 426:555–559.

Du L, Sohr A, Li Y, Roy S. 2022. Gpi-anchored fgf directs cytoneme-mediated bidirectional contacts to regulate its tissue-specific dispersion. Nature Communications. 13:3482.

Duffy JB. 2002. Gal4 system in drosophila: a fly geneticist’s swiss army knife. genesis. 34:1–15.

Edwards KA, Demsky M, Montague RA, Weymouth N, Kiehart DP. 1997. Gfp-moesin illuminates actin cytoskeleton dynamics in living tissue and demonstrates cell shape changes during morphogenesis indrosophila. Developmental biology. 191:103–117.

Fehon RG, Kooh PJ, Rebay I, Regan CL, Xu T et al. 1990. Molecular interactions between the protein products of the neurogenic loci notch and delta, two egf-homologous genes in drosophila. Cell. 61:523–534.

Feinberg EH, VanHoven MK, Bendesky A, Wang G, Fetter RD et al. 2008. Gfp reconstitution across synaptic partners (grasp) defines cell contacts and synapses in living nervous systems. Neuron. 57:353–363.

Furman D, Bukharina T. 2008. How drosophila melanogaster forms its mechanoreceptors. Current genomics. 9:312–323.

Georgiou M, Baum B. 2010. Polarity proteins and rho gtpases cooperate to spatially organise epithelial actin-based protrusions. Journal of cell science. 123:1089–1098.

González-Méndez L, Seijo-Barandiarán I, Guerrero I. 2017. Cytoneme-mediated cell-cell contacts for hedgehog reception. Elife. 6:e24045.

Hadjivasiliou Z, Hunter GL, Baum B. 2016. A new mechanism for spatial pattern formation via lateral and protrusion-mediated lateral signalling. Journal of the Royal Society Interface. 13:20160484.

Hall ET, Dillard ME, Cleverdon ER, Zhang Y, Daly CA et al. 2024. Cytoneme signaling provides essential contributions to mammalian tissue patterning. Cell. 187:276–293.

Hall ET, Dillard ME, Stewart DP, Zhang Y, Wagner B et al. 2021. Cytoneme delivery of sonic hedgehog from ligand-producing cells requires myosin 10 and a dispatched-boc/cdon co-receptor complex. Elife. 10:e61432.

Huang H, Kornberg TB. 2015. Myoblast cytonemes mediate wg signaling from the wing imaginal disc and delta-notch signaling to the air sac primordium. elife. 4:e06114.

Huang H, Kornberg TB. 2016. Cells must express components of the planar cell polarity system and extracellular matrix to support cytonemes. elife. 5:e18979.

Huang H, Liu S, Kornberg TB. 2019. Glutamate signaling at cytoneme synapses. Science. 363:948–955.

Huang J, Zhou W, Dong W, Watson AM, Hong Y. 2009. Directed, efficient, and versatile modifications of the drosophila genome by genomic engineering. Proceedings of the National Academy of Sciences. 106:8284–8289.

Hunter GL, Hadjivasiliou Z, Bonin H, He L, Perrimon N et al. 2016. Coordinated control of notch/delta signalling and cell cycle progression drives lateral inhibition-mediated tissue patterning. Development. 143:2305–2310.

Hunter GL, He L, Perrimon N, Charras G, Giniger E et al. 2019. A role for actomyosin contractility in notch signaling. BMC biology. 17:12.

Jacquemet G, Stubb A, Saup R, Miihkinen M, Kremneva E et al. 2019. Filopodome mapping identifies p130cas as a mechanosensitive regulator of filopodia stability. Current Biology. 29:202–216.

Junyent S, Reeves J, Gentleman E, Habib SJ. 2021. Pluripotency state regulates cytoneme selectivity and self-organization of embryonic stem cells. Journal of Cell Biology. 220:e202005095.

Lacoste J, Soula H, Burg A, Audibert A, Darnat P et al. 2022. A neural progenitor mitotic wave is required for asynchronous axon outgrowth and morphology. Elife. 11:e75746.

Langridge PD, Struhl G. 2017. Epsin-dependent ligand endocytosis activates notch by force. Cell. 171:1383–1396.

Le Borgne R, Schweisguth F. 2003. Notch signaling: endocytosis makes delta signal better. Current biology. 13:R273–R275.

Lewis J. 1996. Neurogenic genes and vertebrate neurogenesis. Current opinion in neurobiology. 6:3–10.

Liu R, Woolner S, Johndrow JE, Metzger D, Flores A et al. 2008. Sisyphus, the drosophila myosin xv homolog, traffics within filopodia transporting key sensory and adhesion cargos. Development. 135.

Liu S, Daly CA, Ogden SK, Zurzolo C. 2026. Regulation and function of specialized membrane protrusions in intercellular communication. Nature Reviews Molecular Cell Biology. pp. 1–18.

Loubéry S, González-Gaitán M. 2014. Monitoring notch/delta endosomal trafficking and signaling in drosophila, In:, Elsevier. volume 534. pp. 301–321.

Mehaffey TM, Hecht CA, White JS, Hutson MS, Page-McCaw A. 2024. Live imaging basement membrane assembly under the pupal notum epithelium. Micropublication Biology. 2024:10–17912.

Melak M, Plessner M, Grosse R. 2017. Actin visualization at a glance. Journal of cell science. 130:525–530.

Miller SW, Rebeiz M, Atanasov JE, Posakony JW. 2014. Neural precursor-specific expression of multiple drosophila genes is driven by dual enhancer modules with overlapping function. Proceedings of the National Academy of Sciences. 111:17194–17199.

Patel A, Wu Y, Han X, Su Y, Maugel T et al. 2022. Cytonemes coordinate asymmetric signaling and organization in the drosophila muscle progenitor niche. Nature communications. 13:1185.

Petruccelli E, Feyder M, Ledru N, Jaques Y, Anderson E et al. 2018. Alcohol activates scabrous-notch to influence associated memories. Neuron. 100:1209–1223.

Powell PA, Wesley C, Spencer S, Cagan RL. 2001. Scabrous complexes with notch to mediate boundary formation. Nature. 409:626–630.

Rambaud B, Joseph M, Tsai FC, De Jamblinne C, Strakhova R et al. 2025. Slik sculpts the plasma membrane into cytonemes to control cell-cell communication. The EMBO journal. 44:2186–2210.

Renaud O, Simpson P. 2001. scabrous modifies epithelial cell adhesion and extends the range of lateral signalling during development of the spaced bristle pattern in drosophila. Developmental biology. 240:361–376.

Renaud O, Simpson P. 2002. Movement of bristle precursors contributes to the spacing pattern in drosophila. Mechanisms of development. 119:201–211.

Roegiers F, Younger-Shepherd S, Jan LY, Jan YN. 2001. Two types of asymmetric divisions in the drosophila sensory organ precursor cell lineage. Nature cell biology. 3:58–67.

Rojas-Rios P, Guerrero I, Gonzalez-Reyes A. 2012. Cytoneme-mediated delivery of hedgehog regulates the expression of bone morphogenetic proteins to maintain germline stem cells in drosophila. PLoS biology. 10:e1001298.

Rosa A, Vlassaks E, Pichaud F, Baum B. 2015. Ect2/pbl acts via rho and polarity proteins to direct the assembly of an isotropic actomyosin cortex upon mitotic entry. Developmental cell. 32:604–616.

Roy S, Huang H, Liu S, Kornberg TB. 2014. Cytoneme-mediated contact-dependent transport of the drosophila decapentaplegic signaling protein. Science. 343:1244624.

Sanders TA, Llagostera E, Barna M. 2013. Specialized filopodia direct long-range transport of shh during vertebrate tissue patterning. Nature. 497:628–632.

Sasaki N, Sasamura T, Ishikawa HO, Kanai M, Ueda R et al. 2007. Polarized exocytosis and transcytosis of notch during its apical localization in drosophila epithelial cells. Genes to Cells. 12:89–103.

Schindelin J, Arganda-Carreras I, Frise E, Kaynig V, Longair M et al. 2012. Fiji: an open-source platform for biological-image analysis. Nature methods. 9:676–682.

Schwabe T, Neuert H, Clandinin TR. 2013. A network of cadherin-mediated interactions polarizes growth cones to determine targeting specificity. Cell. 154:351–364.

Shaya O, Binshtok U, Hersch M, Rivkin D, Weinreb S et al. 2017. Cell-cell contact area affects notch signaling and notch-dependent patterning. Developmental cell. 40:505–511.

Singh H, Brooks E, Jinnai K, Kondo S, Manning SA et al. 2026. The mechanosensitive protein zyxin influences hippo signalling and tissue growth via adherens junctions and basal spot junctions in drosophila. bioRxiv. pp. 2026–02.

Stanganello E, Hagemann AI, Mattes B, Sinner C, Meyen D et al. 2015. Filopodia-based wnt transport during vertebrate tissue patterning. Nature communications. 6:5846.

Stinchfield MJ, Weasner BP, Weasner BM, Zhitomersky D, Kumar JP et al. 2024. Fourth chromosome resource project: a comprehensive resource for genetic analysis in drosophila that includes humanized stocks. Genetics. 226:iyad201.

Sutcliffe C, Nandy N, Revici R, Johnson H, Habib SJ et al. 2025. Ovarian germline stem cell dedifferentiation is cytoneme dependent. Proceedings of the National Academy of Sciences. 122:e2426145122.

Troost T, Schneider M, Klein T. 2015. A re-examination of the selection of the sensory organ precursor of the bristle sensilla of drosophila melanogaster. PLoS genetics. 11:e1004911.

Tseng Q, Duchemin-Pelletier E, Deshiere A, Balland M, Guillou H et al. 2012. Spatial organization of the extracellular matrix regulates cell–cell junction positioning. Proceedings of the National Academy of Sciences. 109:1506–1511.

Uemura T, Oda H, Kraut R, Hayashi S, Kotaoka Y et al. 1996. Zygotic drosophila e-cadherin expression is required for processes of dynamic epithelial cell rearrangement in the drosophila embryo. Genes & development. 10:659–671.

Vandervorst P, Ghysen A. 1980. Genetic control of sensory connections in drosophila. Nature. 286:65–67.

Yoshigi M, Hoffman LM, Jensen CC, Yost HJ, Beckerle MC. 2005. Mechanical force mobilizes zyxin from focal adhesions to actin filaments and regulates cytoskeletal reinforcement. The Journal of cell biology. 171:209–215.

Zhang C, Brunt L, Ono Y, Rogers S, Scholpp S. 2024. Cytoneme-mediated transport of active wnt5b–ror2 complexes in zebrafish. Nature. 625:126–133.

